# Studies on the International Space Station to assess the effects of microgravity on iPSC-derived neural organoids

**DOI:** 10.1101/2023.08.10.552814

**Authors:** Davide Marotta, Laraib Ijaz, Lilianne Barbar, Madhura Nijsure, Jason Stein, Twyman Clements, Jana Stoudemire, Paula Grisanti, Scott A. Noggle, Jeanne F. Loring, Valentina Fossati

## Abstract

Exposure to microgravity in low-Earth orbit (LEO) has been shown to affect human cardiovascular, musculoskeletal, and immune systems. Post-flight brain imaging indicates that reports about astronauts and mouse models suggest that microgravity may cause intracranial fluid shifts and possibly alter white and gray matter of the brain [1]. To focus on the effects of microgravity on the brain, we used induced pluripotent stem cells (iPSCs) to produce three-dimensional (3D) human neural organoids as models of the nervous system. We studied iPSCs derived from four individuals, including people with the neurological diseases primary progressive multiple sclerosis (PPMS) and Parkinson’s disease (PD) and non-symptomatic controls. We patterned the organoids toward cortical and dopaminergic fates representing regions of the brain affected by MS and PD, respectively. Microglia were generated from the same cell lines and integrated into a portion of the organoids. The organoids were maintained for 30 days in a novel static culture system on the International Space Station (ISS) and live samples were returned to Earth. The post-flight samples were evaluated using histology, transcriptome and secretome analysis. Microglia-specific genes and secreted proteins were detectable in the microglia-containing organoid cultures. The gene expression analyses of individual organoids cultured in LEO and on Earth suggest that cell proliferation was lower and neural cells were more mature in samples that were cultured in LEO. These experiments lay the groundwork for further studies, including long term studies to investigate the effects of microgravity on the brain. With two more missions using similar cells, we are determining whether this effect of microgravity is consistent in separate experiments. Such studies may ultimately aid in developing countermeasures for the effects of microgravity on the nervous systems of astronauts during space exploration and suggest novel therapeutic interventions for neurological diseases on Earth.

## Introduction

Experimental biological research on the International Space Station (ISS) in low-Earth orbit (LEO) is intended to both understand the effects of microgravity on human health and to investigate the use of the ISS as a platform for improvements in disease modeling and drug development. The studies performed so far have indicated that microgravity causes changes in musculoskeletal systems and the peripheral immune system, and can affect vestibular function and cognition. Cell types derived from human induced pluripotent stem cells (iPSCs) can be used to build relevant *in vitro* models of human organs to study functional and structural changes that may be important to human health in space and on Earth. Because of our interest in neurodegenerative disease, we included iPSC-derived neural cells from individuals with primary progressive multiple sclerosis (PPMS) and Parkinson’s disease (PD). We created three-dimensional aggregates of cells, known as organoids, that contained iPSC-derived cortical neurons to model PPMS or dopaminergic neurons to model PD. To improve the utility of these simple neuronal models, we added an additional brain-resident cell type, microglia, which are components of the brain’s immune system, to a portion of the organoids.

We developed new protocols specifically for the experiments on the ISS, including a novel cell culture system, transport methods, and post-flight recovery and analysis. We generated the organoids in our lab, transported them to the Space Station Processing Facility (SSPF) at Kennedy Space Center (KSC), and placed each organoid in a cryovial containing 1 ml of culture medium (see Methods); the vials were loaded into flight hardware that maintained temperature control during transport to and return from the ISS. Passive dosimeters were included in the flight hardware and station radiation monitoring logs were used to measure radiation levels during the mission. The cultures were launched to the ISS on the SpaceX 19^th^ Commercial Resupply Services mission for NASA (SpX CRS-19) and maintained in static culture onboard the ISS for 30 days, followed by live return for post-flight analysis (**Fig. 1**). We maintained a parallel ground control experiment at the KSC laboratory.

**Figure 1.**
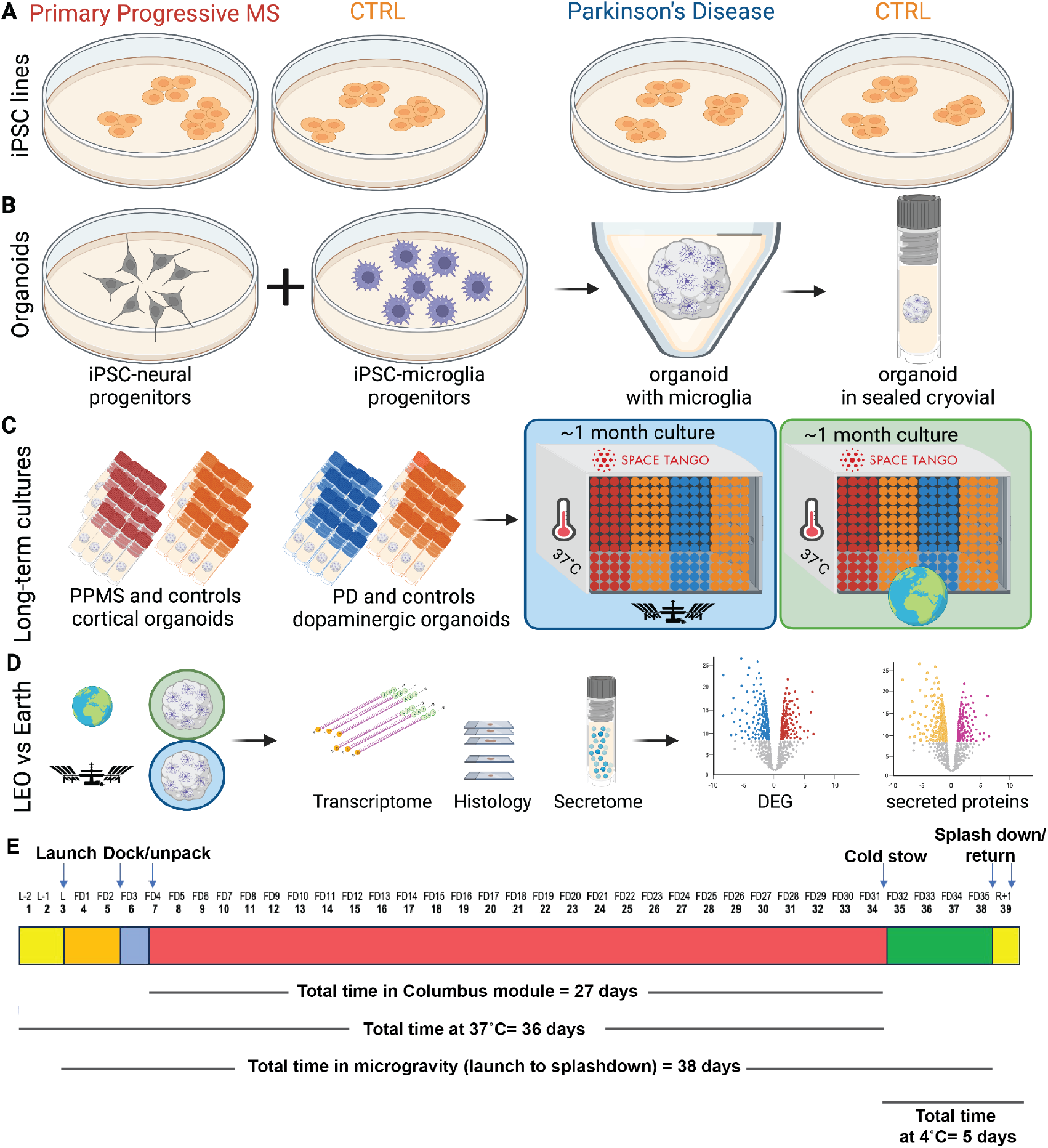
Experimental design. **A**. iPSCs from four individuals were selected for experiments, two controls and two with neurodegenerative disease. **B**. iPSCs were differentiated into cortical or dopaminergic neuron precursors, aggregated to form organoids, and matched microglia derived from each of the cell lines were added to half of the organoids. Each organoid was placed in a separate cryovial with 1 ml of culture medium and sealed for the duration of the experiment. One set of tubes was transported to the ISS on the CRS-19 mission. **C**. Matching sets of cryotube cultures were incubated for a month at 37°C on ground and onboard the ISS. **D**. Upon return to Earth, both sets of organoid cultures were analyzed by multiple methods. **E**. Timeline of the mission.

The ground and LEO samples were analyzed by multiple methods: post-flight viability, immunohistochemistry, whole genome RNA sequencing, and analysis of proteins secreted into the culture medium. The returning cells were alive and extended neurites when placed on culture dishes. We observed significant differences in gene expression and protein secretion associated with LEO samples in both cortical and dopaminergic organoids, suggesting changes in cell-cell signaling. The radiation monitoring and gene expression analysis indicate that microgravity, not radiation, was responsible for these differences. This pioneering experiment demonstrates that complex cultures of iPSC-derived central nervous system (CNS) cells can be successfully maintained in LEO for long periods of time, establishing the foundation for future studies. Our preliminary findings from transcriptomic and secretome analyses support the hypothesis that microgravity affects CNS cells and warrants additional investigations.

## Materials and Methods

### Cell lines

This study was performed using four iPSC lines (**Table 1**). iPSC line 051121-01-MR-017 and AK003-01-MR-008 were previously reprogrammed from dermal fibroblasts with an mRNA/miRNA method [2]. iPSC line HDF410iPS504 [3] and iPSC line UEC741iPS517 were previously reprogrammed from dermal fibroblasts and urine epithelial cells respectively using non-integrating Sendai virus [4]. iPSC lines were expanded and maintained in mTeSR1 medium, following published methods [5], assessed for pluripotency using RNAseq and PluriTest [6] and for genomic integrity with single nucleotide polymorphism (SNP) microarrays [7].

**Table 1:**
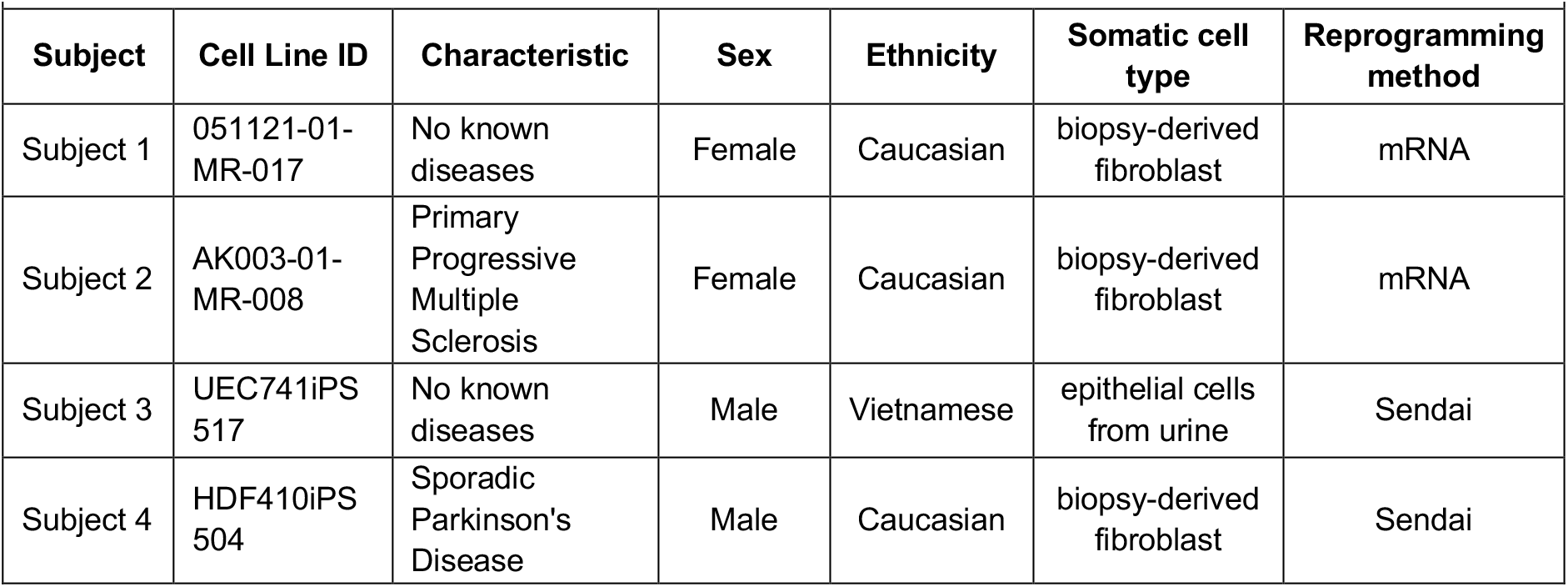
Cell lines and characteristics.

### Differentiation and assembly of organoids and microglia

**Figure 1 and Supplemental Figure 2** illustrate the steps involved in the experiment from culture of the organoids to post-flight recovery and analysis. iPSC line 051121-01-MR-017 and AK003-01-MR-008 were differentiated into cortical neural precursors using established protocols [8] and iPSC line HDF410iPS504 and iPSC line UEC741iPS517 were differentiated into dopaminergic neural precursors [3]. Twenty-five days after differentiation was initiated, both cortical and dopaminergic precursors were dissociated and cryopreserved. Isogenic microglia were derived from all of the cell lines [9]. A week before the estimated launch date, cryopreserved neural precursors were thawed, washed in DMEM/F12, and spun at 300g for 5 min. Cell pellets were resuspended in cortical neuron medium [8] or dopaminergic neuron medium [3] and were plated at a density of 1×10^5^ cells/well onto ultra-low attachment (ULA) 96-well V bottom plates (PrimeSurface® 3D culture spheroid plates (ULA plates, Sbio MS-9096VZ) and incubated overnight at 37°C, 5% CO_2_ to allow cell aggregation into organoids. For half of the samples, isogenic iPSC-derived microglia were added to the neural organoids after 24 hours, and a medium exchange was performed, adding IL-34 (100ng/mL) and GM-CSF (10ng/mL) to promote microglia maturation. Cultures were then incubated for an additional 24 hours before being shipped at 4°C to the Kennedy Space Center for payload preparation and launch.

### Payload preparation, launch, and splashdown

Organoids were individually transferred into a cryovial (NUNC™ Coded Cryobank Vial, Thermo Fisher Scientific #374088), containing 1 mL of cortical or dopaminergic medium with the addition of IL-34 (100ng/mL) and GM-CSF (10ng/mL) for those containing microglia, in DMEM/F-12 medium buffered with 15mM HEPES (Thermo Fisher Scientific #11330). The cryovials were closed and sealed with parafilm and loaded into a CubeLab, a miniaturized incubator that maintains a stable temperature of 37°C. The CubeLab was then transferred to the rocket and after two days, launched into low-Earth orbit onboard the SpaceX 19th Commercial Resupply Services mission (SpX CRS-19) to the ISS for a 30-day stay in microgravity. Under similar conditions, ground control samples were kept at the Kennedy Space Center. After 30 days, the Dragon capsule carrying the organoids back to Earth splashed down in the Pacific Ocean, and the CubeLab was opened six days post undock from the ISS (see **Fig. 1**).

### Post-flight culture of organoids

As soon as the CubeLab was opened, the viability of the organoids was assessed by plating 4 organoids separately on culture dishes coated with laminin/fibronectin and cultured for several more days at 37o C in differentiation medium conventionally buffered for CO_2_.

### RNA extraction and sequencing

**Table S2** lists the individual organoids assayed by RNA sequencing. Twenty organoids cultured in LEO and 22 from ground control, both with and without added microglia, were snap-frozen and stored at -80°C. RNA was extracted using the Qiagen RNeasy Micro kit (QIAGEN #74004). To maximize the yield, the RNA was eluted into 12μL of ultrapure DI water. RNA quality and concentration was assessed on Bioanalyzer Pico with RIN reported (**Table S2**). The RNA was sequenced by an outside provider (Novogene; https://www.novogene.com/us-en/), which used an ultra-low input preparation of samples with a minimum of 10ng of RNA and the Illumina NovaSeq Platform for high-throughput PE150 sequencing. A pilot subset of samples was cDNA amplified before sequencing with the Clontech SMART kit employing Clontech SMART CDS Primer II A. Raw FASTQ data were filtered for adapters with fastp v0.23.0 software and aligned to human genome using STAR v 2.7.10a with indexes generated from the Homo_sapiens GRCh38 primary assembly and the GRCh38.107.gtf build ensemble annotations. The featureCounts v2.0.3 software was used to generate a counts matrix for differential expression analysis. The organoids were analyzed individually, and after datasets were filtered, data from organoids composed of the same cell types were combined to increase the power of the analysis.

### Differential gene expression, Gene Set Enrichment Analysis (GSEA), Transcription Factor Enrichment Analysis (TFEA)

The RNA sequencing data were analyzed using the R packages DESeq2 [10], and Gene Set Enrichment Analysis (GSEA) GO term analysis analyzed using gProfiler [11] and ggplot2 [12]. TFEA [13] was performed on gene sets that were differentially expressed in LEO compared to the parallel samples on the ground.

### Analysis of secreted proteins using Proximity Extension Assay (PEA)

Culture medium from vials cultured in LEO or on ground (**Table S5**) was removed and frozen at -20°C. Proteomic analysis was performed by Olink (Olink Proteomics AB, Uppsala, Sweden) using the Olink® Explore 1536 Olink protocol technology based on the Proximity Extension Assay (PEA) [14]. Olink® Explore 1536 included a combination of 4 separate Olink® Explore 384 panels: Olink® Explore 384 Inflammation (Product Code 91101), Olink® Explore 384 Oncology (Product Code 91102), Olink® Explore 384 Cardiometabolic (Product Code 91103), and Olink® Explore 384 Neurology (Product Code 91104). Each panel used 384 pairs of antibodies conjugated to unique DNA sequences that hybridize and amplify upon binding to their target proteins. The resulting DNA amplicons were then quantified by sequencing using the Illumina® NovaSeq system. The output data were given as Normalized Protein eXpression (NPX) values, which are log2-transformed relative quantification values derived from optimized standard curves for each assay that are normalized for inter-plate variation and internal controls. Each sample was assessed for its overall quality based on the number of detected proteins and the percentage of missing values. Samples with fewer than 46% detected proteins or more than 25% missing values were excluded from further analysis. The secretome data were analyzed using the OlinkAnalyze R package v3.4.1. As with the RNAseq data, we analyzed cortical organoids and dopaminergic organoids separately. We used the olink_lmer function to obtain estimated means for each assay, fitting a linear mixed model with NPX as the dependent variable, microglia presence and ground vs. LEO treatment condition as the fixed effect, and cell line as random effect. Post-hoc analysis with olink_lmer_posthoc was used to obtain relative NPX expression estimates for the ground vs LEO treatment conditions for the cortical and dopaminergic organoids.

### Staining, image acquisition, and processing

The live organoids from LEO and ground controls were fixed with a 4% paraformaldehyde (PFA) solution in PBS for 30 minutes. The organoids were stained as whole mounts [15] using an antibody to MAP2 (Ab5392, dilution 1:1000) to detect neurites and Hoechst dye for labeling nuclei. The labeled organoids were imaged using a ZEISS confocal microscope (LSM780), and images were processed and edited using Imaris software (Oxford Instruments, Abingdon, UK).

### Radiation monitoring

Radiation Environmental Monitoring (REM) and Hybrid Electronic Radiation Assessor (HERA) devices are currently operational on the ISS [16]. Station monitoring logs adjacent to the payload storage location on the ISS were provided for post-flight evaluation as a method for measurement of radiation exposure during the mission. The SpX CRS-19 payload was stowed in the European Space Agency (ESA) Columbus Laboratory during the missions. Timestamp-associated REM measurements from radiation logs provided by NASA for this location are reported as daily GCR (mGy/d) and SAA (mGy/d) doses and combined to report averaged daily exposures. Cumulative exposures were calculated for the period defined as the time from docking of the Dragon vehicle to the ISS through the day of vehicle undocking.

## Results

### Culture of brain organoids in microgravity

As a model to investigate the effects of microgravity on the brain, we developed methods to maintain human iPSC-derived neural organoids in low-Earth orbit (LEO) onboard the International Space Station (ISS). We differentiated cortical and dopaminergic neurons from four different donors (**Table 1**). We chose cell lines that are in use in our laboratories for other projects studying MS and PD. Since the affected individuals had sporadic neurogenerative disease without a known genetic basis, our studies were not powered to detect disease associated differences among the iPSCs from MS and PD and the controls. However, we hypothesized that we might detect differences associated with culture of the organoids in LEO. Since neurons originate from ectoderm, and microglia from extraembryonic yolk sac, during the early stages of embryogenesis, we derived microglia from each of the iPSC lines separately and added the matching cells to half of the organoids (**Fig. 1**). Two sets of organoids were cultured in LEO or on Earth for approximately one month in 1ml of medium in cryovials without medium changes. The organoids increased in size both in LEO and on ground (**Fig. 2A**). Upon their return to Earth, several of the organoids were transferred to culture dishes, where they attached to the surface and rapidly elaborated networks of neurites (**Fig. 2B**). The cortical organoids showed the typical neural rosettes of this neuronal subtype (**Fig. 2C, Fig. S1**).

**Figure 2.**
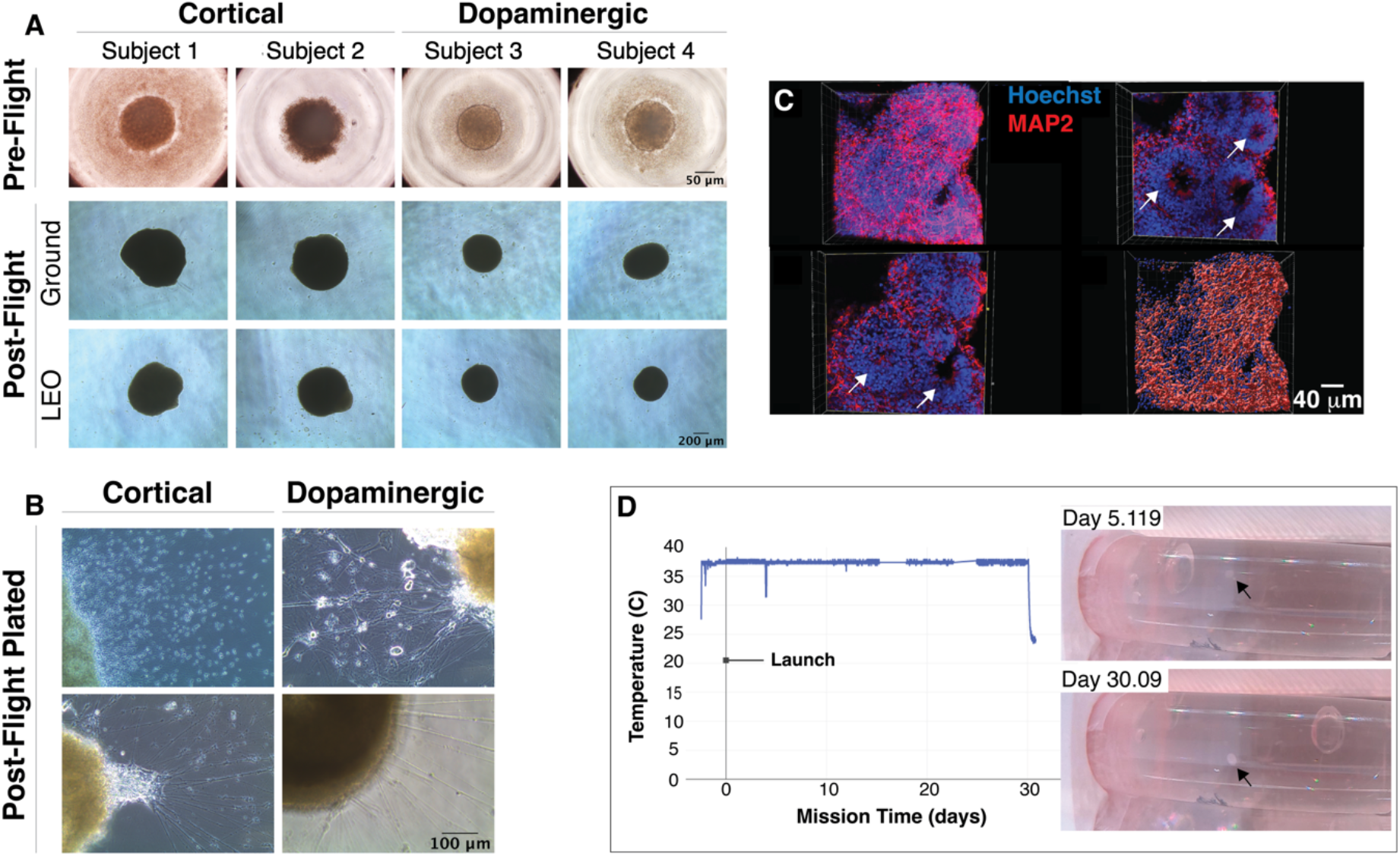
Neural organoids cultured in static systems without media exchange are viable after 30 days. **A**. Cortical and dopaminergic neural organoids, from pre-flight stage to the post-flight stage in low-Earth orbit (LEO) and on ground. Organoids cultured post-flight (**B)** show outgrowth of neural projection and radial glia after 30 days culture in low-Earth orbit (LEO). The cortical organoid in **C** shows neural rosettes (Hoechst) and neurites (MAP2). The telemetry in **D** shows the constant temperature maintained inside the CubeLab onboard the ISS and the images show the organoid (arrow) cultured in static systems at mission day 5.119 and day 30.09 (images taken onboard the ISS).

Monitors in the incubator showed that the temperature remained constant at 37° C, then was lowered to 4° C for the return to Earth, and the organoids could be visualized in the cryotubes during their time on the ISS (**Fig. 2D**).These data show the viability of the CNS organoids cultured in static systems without media exchange during long-term cultures in LEO and on ground.

### Gene expression profiles of organoids are altered in LEO

**Figure 3** illustrates the results of global gene expression analysis. **Table S1** lists the individual organoids that were analyzed by RNAseq. After quality control and filtering, principal component analysis (PCA) of all organoids demonstrated a clear distinction based on organoid type (dopaminergic vs. cortical organoids) and the cell line used to generate the organoids. These variables accounted for the majority of the data’s variance. However, distinctions based on ground vs. LEO conditions were also identified by PCA and clustering analysis (**Fig. 3**). To investigate the effects of LEO on each organoid type, we performed separate analyses for each organoid type (cortical or dopaminergic) for all cell lines. After controlling for the sequencing library preparation method, organoid type, cell line, and the addition of microglia to the organoids, we conducted a differential gene expression analysis between organoids cultured on Earth (ground condition) and those cultured in LEO on the ISS. For the cortical organoids, we found 4,849 differentially expressed genes (DEGs) between LEO and ground conditions (2,369 higher in LEO vs. 2,480 lower in LEO, at adjusted p-value < 0.05), and 6,385 DEGs for the dopaminergic organoids (2,911 higher in LEO vs. 3,474 lower in LEO, at adjusted p-value < 0.05) (**Table S2**).

**Figure 3.**
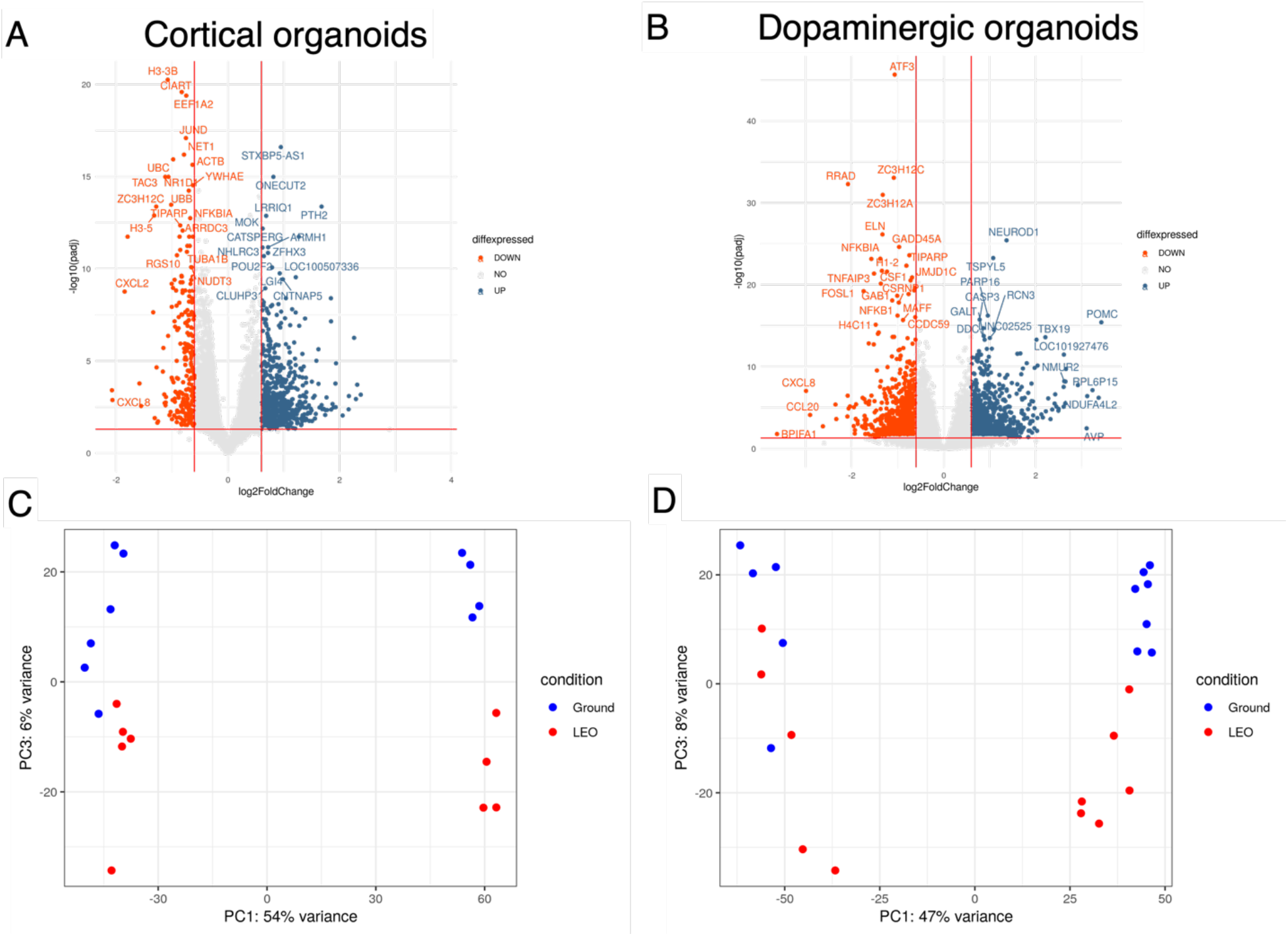
Microgravity alters gene expression in neural organoids. Differentially expressed genes (DEGs) in **A** and **B** were analyzed using DESeq2 and the volcano plot created using the packages ggplot2 and ggrepel. The log_2_ fold change indicates the mean expression for each gene. Genes with significant enrichment (padj <0.05) are shown as red (lower in LEO, “down”) or blue (higher in LEO, “up”) across replicates (log_2_ fold change >1 or < -1, respectively). The volcano plots show the differentially expressed genes (DEGs) – low-Earth orbit (LEO) versus ground – of the iPSC-derived cortical organoids (**A**), and the iPSC-derived dopaminergic organoids (**B**). ND = no difference. **C**. PCA plot showing the differences between LEO (red) and ground (blue) of the cortical organoids from 2 iPSC lines. **D**. PCA plot showing the differences between LEO (red) and ground (blue) of the dopaminergic organoids from 2 iPSC lines.

To understand the biological processes impacted by microgravity, we performed Gene Set Enrichment Analysis (GSEA) and Gene Ontology (GO) analysis on the ground vs. LEO DEGs. The highest-ranking differentially expressed gene sets were linked to *DNA repair, mitotic checkpoint*, and *maturation* in LEO (**Table S3**). In the combined analysis, several terms were significantly represented for cortical organoids, including terms associated with ciliated cells (*axonemal microtubule*: EFHB, CFAP210, PIERCE1, Padj=2.799×10-2), and glial differentiation (*positive regulation of gliogenesis*: HES1, CXCR4, LRP2, Padj=3.59×10-2). In dopaminergic organoids in LEO, we observed an enrichment of terms related to neuronal differentiation and maturation, including *synaptic transmission, cholinergic* (CHRNA5, CHRNB3, CHRNE, SLC5A7, SLC18A3, Padj=1.238×10-2), *nicotine effect on dopaminergic neurons* (DDC, CHRNA5, DRD2, Padj=2.98×10-2), and *acetylcholine-gated channel complex* (CHRNA5, CHRNB3, CHRNE, Padj=4.645×10-2). In dopaminergic organoids, we also observed enriched terms related to hypothalamus development, including *corticotropin-releasing hormone signaling pathway* (CASP3, POMC, TBX19, Padj=1.703×10-2) and *growth hormone secretion* (DRD2, GHSR, GHRH, Padj=2.46×10-2). We also noted an enrichment for genes and terms related to Wnt signaling and *diencephalon development* in both organoid types (**Table S3**).

Genes related to stress and apoptosis were commonly found to be lower in LEO for both organoid types, including those associated with *apoptosis, cellular responses to stress*, and *cellular senescence* (**Table S3**). Inflammatory pathway markers and terms were also globally generally lower in LEO. These include genes associated with *Epstein-Barr virus infection, cell chemotaxis, chemokine receptor binding*, and the *IL-18 signaling pathway*. Further, *oxidative damage response genes* were also found to be lower in LEO than on Earth. Transcription factor enrichment analysis (TFEA) of the DEGs highlighted the common transcription factor binding sites of sets of genes (**Table S4**). This analysis revealed co-regulated sets of genes associated with neural differentiation, and confirmed the dominant involvement of Wnt signaling pathways associated with neural development that are underlying the differences between organoids in LEO and on the ground. These results, increases in differentiation-associated markers and decreases in proliferation-associated genes, are consistent with the idea that neuronal maturation is enhanced in LEO compared to ground controls.

### Microgravity influences protein secretion

To assess the effects of microgravity on secretion of proteins, culture medium was collected from vials containing single organoids and analyzed using Olink® Explore 1536. We used Olink® Explore 1536, a high-throughput proteomics platform based on the Proximity Extension Assay (PEA) technology, to measure the expression levels of 1,536 human proteins in 30-day conditioned supernatants from ground and LEO organoid samples. Overall, we found NPX levels lower in LEO vs ground. However, both of the organoid types showed LEO vs. ground differentially secreted proteins (DSPs) that were either upregulated or downregulated (**Table S5**). For example, in cortical organoids, SFRP1, a modulator of the Wnt signaling pathway, was lower in LEO, whereas NTF3, a neurotrophic factor, is upregulated in LEO. Multiple microglia markers, including CXCL8 and CD14, were found to be consistently upregulated in the samples that had been loaded with microglia (**Fig. 4A**). Additionally, we found several proteins were upregulated in LEO but only in the presence of microglia, including the inflammatory mediator IL1R1. (**Fig. 4B** and **Table S5)**. Together these data suggest complex alterations in differentiation and inflammatory pathways in response to the microgravity environment in LEO.

**Figure 4.**
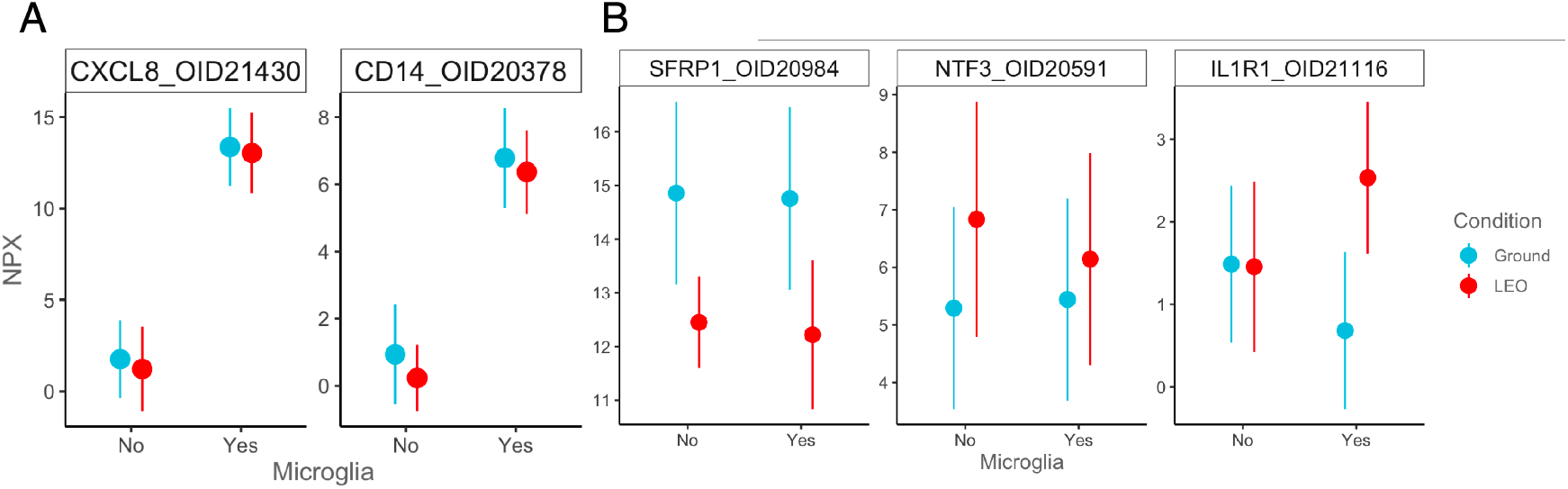
Microgravity affects the secretome of organoids cultured on the ISS. **(A)**. Markers for microglia (CXCL8 and CD14) could be detected in supernatants from organoids with microglia added. (B) Examples of significantly differentially detected proteins in LEO (red) vs. ground (turquoise) showing examples of protein SFRP1 overall decreased in LEO, NTF3 overall increased in LEO, and IL1R1 only increased in LEO in organoids containing microglia.

### Radiation exposure of organoids onboard the ISS

The radiation field inside the ISS fluctuates due to different local shielding thicknesses and the orbital changes of the ISS. The cultures were located in the European Space Agency (ESA) Columbus Laboratory. Timestamp-associated REM measurements from radiation logs provided by NASA for this location are reported as the amount of radiation (mGy/d) daily Galactic Cosmic Radiation (GCR) and Southern Atlantic Anomaly (SAA) doses and combined to report average daily exposures. Cumulative exposures were calculated for the period defined as the time period from docking of the Dragon vehicle to the ISS through the day of vehicle undocking. The station radiation logs provided by NASA for this location reported GCR daily average dose of 0.153 mGy (milligrays) per day and a SAA daily average dose of 0.231 mGy/d, for a combined daily radiation exposure dose of 0.384 mGy/d. Cumulative radiation exposure using this combined daily average dose would be 11.5 mGy (approximately 12 mGy) over the 30 days the payload was stowed in the Columbus module. The calculation of radiation exposure, measured in millisieverts (mSV), is dependent on many variables, including the amount of shielding of the object and tissue-specific radiation sensitivity. While the organoid-specific exposure factors relative to whole body exposure have not been determined, **Table 2** provides a list of approximate radiation exposures on Earth and on the ISS.

**Table 2.** Comparison of radiation dosage levels.

| **Table 2. Comparison of radiation dosage levels** |  |  |
| --- | --- | --- |
| **Location** | **Estimated Radiation** | **Source** |
| Earth (annual total) | Total 3 - 4 mSv/year | Natural sources (rocks, cosmic rays penetrating atmosphere) |
| Pilot (long haul Polar routes; annual total) | Total 6 mSv/year | Cosmic radiation |
| Astronaut on ISS (daily; 6-month mission) | 0.5 mSv/day<br>Total 90 mSv/6 months | GCR/SAA (Galactic Cosmic Radiation (GCR) / Southern Atlantic Anomaly (SAA)) |
| Payload SpX - 19 (30-day mission) | 0.4 mSv/day<br>Total 12 mSv/30 days | GCR/SAA |
| NASA threshold for astronauts (lifetime) | Total 600 mSv/lifetime | GCR/SAA |
| Russian & Canadian threshold for astronauts (lifetime) | Total 1000 mSv/lifetime | GCR/SAA |

On Earth, humans are exposed to 3 to 4 mSV of radiation per year, mostly from natural sources including some types of rocks and a few cosmic rays that get through the atmosphere. At higher altitudes and latitudes passengers are exposed to elevated, although insignificant, levels of cosmic radiation. Crew and pilots serving on long-haul polar routes however may receive an annual effective dose of up to 6 mSv [17].

We consider the estimated 12 mGy of cumulative dose over the 30-day mission to be indicative of relatively low exposure levels during the time of the mission. While it is unlikely that there could have been acute exposure during the 5 days between launch and when the payload was transferred to the Columbus module, the remaining 30 days at 37°C would be likely to be representative of consistent effects. While cumulative radiation exposure on long-duration space missions is an area of intensive study and interest for NASA, and the subject of radiation exposure to cells cultured on the ISS warrants further investigation, the limited exposure of payloads during short duration missions is more similar to risks posed by common radiological procedures (**Table S6**), suggesting the differences observed between flight and ground control samples are more likely related to the microgravity effect than exposure to radiation on the ISS during these missions.

## Discussion

We report here the results of experiments comparing human neural organoids cultured on the ISS with those cultured on Earth. The spherical organoids, about 500 μm in diameter, were formed in microwells on the ground, then transferred to individual cryovials containing 1 ml of culture medium, which is more than 10,000 times the volume of an organoid. After 30 days in the sealed vials, the organoids remained healthy, and when plated onto adherent surfaces, rapidly extended dense webs of neurites.

The organoids were designed to be simplified representations of different parts of the brain, containing either iPSC-derived cortical or dopaminergic neurons with or without incorporation of small populations of matched microglia derived separately from the same cell lines. We chose these two neuronal cell types because they are affected by different neurodegenerative diseases. Cortical neurons are damaged in multiple sclerosis (MS), and dopamine neurons are lost in Parkinson’s disease (PD). We included iPSCs from two individuals for each cell type: cortical neurons were derived from iPSCs from a person with primary progressive MS and a control, and dopamine neurons from a person with sporadic PD and a different non-diseased control. We incorporated microglia, which are one of the immune cell types in the brain, in order to monitor possible immunological effects of long-term culture in microgravity. Microglia had to be derived separately from the same iPSC lines as the neurons because they do not originate in the neural ectoderm of the brain. They are derived from the extraembryonic yolk sac of the developing embryo and migrate into the brain. We were able to detect specific markers of microglia among the proteins secreted into the culture medium from the microglia-containing organoids over the course of incubation.

Individual organoids were analyzed separately by RNA sequencing; the main differences were associated with the specific cell line and RNA preparation method. When these differences were filtered out, the profiles were very similar within the groups of the same neuronal cell type. Remarkably, all of the types of organoids showed significant differences depending on whether they were cultured in LEO or on Earth. Since this was the first time that cultures like these had been examined in LEO, we did not know what to expect, but were anticipating changes that reflected distress in the LEO samples. Instead, we found little evidence of stress, inflammation or apoptosis, and no signs of hypoxia in either group. Interestingly, the organoids cultured on Earth, not the LEO samples, had slightly higher levels of stress-related genes.

The main differences in gene expression pointed toward greater maturity and decreased proliferation in the LEO cultures. This difference is consistent with the observed differences in elements of Wnt signaling pathways. The Wnt family of secreted proteins is highly conserved among species and is integral to cell-cell communication during embryonic development. Wnt signaling is highly complex and controls cell proliferation, migration, and differentiation, especially in the brain [18].

Since there is no precedent for these experiments, the mechanism of this phenotypic change is not yet clear. Previous efforts to accelerate human neuronal differentiation *in vitro* have required genetic interventions or chemical modification of culture conditions, but our ground-based and LEO cultures were not manipulated in this way. There are clues to the possible mechanisms of the observed maturation acceleration, in experiments designed to understand the species-specific time course of neural differentiation [19]. While the sequence of events that occur in brain development is conserved across species, the time scales differ widely, and even when human neural precursors are transplanted to mouse brain, they adhere to their intrinsic time scale. Similarly, *in vitro*, the species is the dominant factor in the timing of neural differentiation. The anomaly that we observed is likely due to microgravity rather than other possible factors, such as radiation exposure; the level of radiation exposure is relatively low and comparable to the exposure levels of astronauts over the same time period.

This is the first report of the effects of microgravity using an organoid model of the central nervous system. With this mission, we established a reproducible and relevant model that revealed intriguing microgravity-induced changes, unexpectedly finding that culture in LEO resulted in more mature, less proliferative neuronal cells. To investigate the mechanisms underlying microgravity’s effect on the neural organoids, we have completed two more experiments on the ISS, launched on CRS-24 and CRS-25, and have additional missions planned for 2024. With each flight, we are adding more complexity to the experiments, including additional cell lines, some with genetic forms of neurodegenerative disease, additional neuronal cell types, and more sophisticated methods of analysis. We hope to use the perturbation of microgravity to better understand neurological disorders. The data reported here form the basis for further explorations, and our goal is to make use of the opportunity to study neural cells in low Earth orbit to understand neurodegenerative disease and ameliorate potentially adverse neurological effects of space travel.

## Acknowledgments

This work was supported by the National Stem Cell Foundation and ISS National Laboratory. We thank Andres Bratt-Leal for his help in the early stages of this project, Olink for assistance with secretome analysis, Jennifer Fogarty for discussions on radiation data, and Karl Willert for his help in interpreting Wnt signaling pathways. Figure 1 was created using Biorender.com.

## Notes

### Competing Interest Statement

The authors have declared no competing interest.

