## Supplemental Figures and Tables for "Studies on the International Space Station to assess the effects of microgravity on iPSC-derived neural organoids"

**Supplementary materials**

**Methods**

1. Details about the parameters of gene expression analysis.

**Figures**

1. S1: Microglia incorporate into organoids
2. S2: Differentiation Protocols

**Tables**

**S1:** List of individual organoids
**S2**: List of all DEGs – excel
**S3**: List of GO terms and GSEA
**S4**: TFEA list- all transcription factor binding sites for all conditions
**S5**: List of data from o-link
**S6**: List of radiation exposures for medical procedures

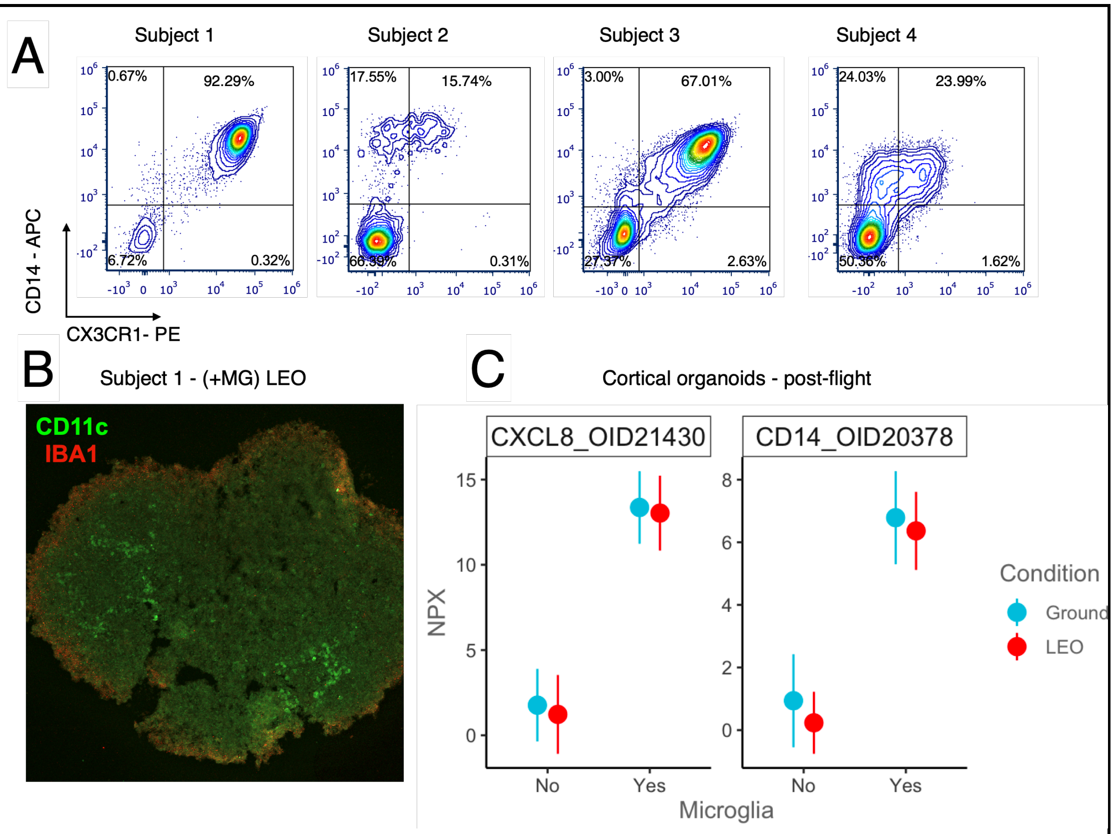


**Figure S1. Microglia incorporate into organoids. (A)** Pre-flight check for microglia markers CD14 and CX3CR1 by FACS analysis. **(B)** Post-flight immunofluorescent stain for CD11c (green) and IBA1 (red) for a cortical organoid from Subject 1 shows incorporation of microglia throughout the organoid. **(C)** Post-flight Olink analysis of culture supernatant from all cortical organoids for microglia markers CXCL8 and CD14 are consistently significantly higher in samples with microglia added.

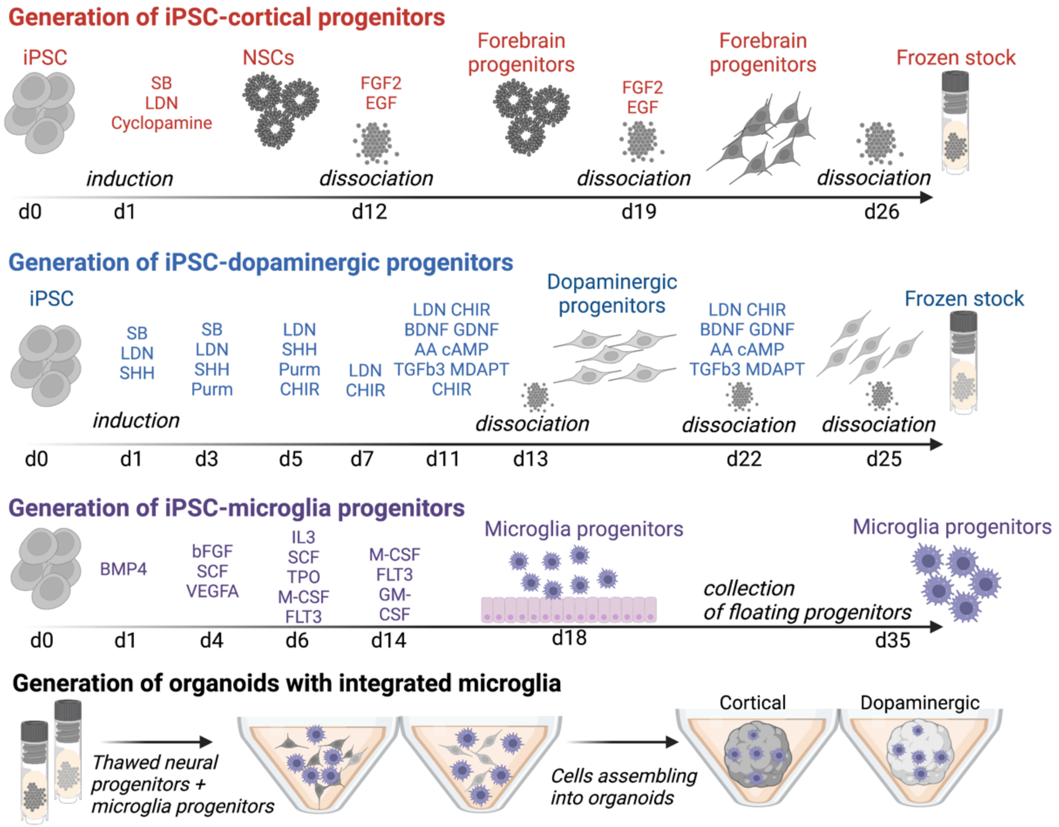


**Figure S2. Differentiation protocols for neural and microglia progenitors and assembly of organoids.** Schematic for differentiation protocols used to generate cortical and dopaminergic neural progenitors and microglia progenitors used to assemble organoids with integrated microglia.

**Supplementary Table 1. Samples analyzed by RNAseq**

| Sample reference ID | Ground/LEO | Subject ID | Cell Line ID | Microglia | Library prep | RNA RIN |
|---|---|---|---|---|---|---|
| **Cortical organoids** | | | | | | |
| b31 | LEO | Subject1 | 051121-01-MR-017 | - | Nextera | 8.3 |
| b33 | LEO | Subject1 | 051121-01-MR-017 | - | Nextera | 7.6 |
| b39 | LEO | Subject1 | 051121-01-MR-017 | + | Nextera | 7.4 |
| b40 | LEO | Subject1 | 051121-01-MR-017 | + | Pilot-TruSeq | 7.8 |
| b41 | LEO | Subject1 | 051121-01-MR-017 | + | Nextera | 8.9 |
| b44 | LEO | Subject2 | AK003-01-MR-008 | - | Nextera | 7.6 |
| b45 | LEO | Subject2 | AK003-01-MR-008 | - | Nextera | 7.7 |
| b50 | LEO | Subject2 | AK003-01-MR-008 | + | Nextera | 7.5 |
| b51 | LEO | Subject2 | AK003-01-MR-008 | + | Nextera | 7.9 |
| a29 | Ground | Subject1 | 051121-01-MR-017 | - | Nextera | 6.6 |
| a30 | Ground | Subject1 | 051121-01-MR-017 | - | Pilot-TruSeq | 6.4 |
| a31 | Ground | Subject1 | 051121-01-MR-017 | - | Nextera | 7.1 |
| a36 | Ground | Subject1 | 051121-01-MR-017 | + | Pilot-TruSeq | 7.0 |
| a37 | Ground | Subject1 | 051121-01-MR-017 | + | Nextera | 6.9 |
| a38 | Ground | Subject1 | 051121-01-MR-017 | + | Nextera | 7.1 |
| a42 | Ground | Subject2 | AK003-01-MR-008 | - | Nextera | 7.2 |
| a43 | Ground | Subject2 | AK003-01-MR-008 | - | Nextera | 7.7 |
| a48 | Ground | Subject2 | AK003-01-MR-008 | + | Nextera | 7.4 |
| a49 | Ground | Subject2 | AK003-01-MR-008 | + | Nextera | 7.3 |
| **Dopaminergic organoids** | | | | | | |
| b1 | LEO | Subject3 | UEC741iPS517 | - | Nextera | 7.8 |
| b2 | LEO | Subject3 | UEC741iPS517 | - | Pilot-TruSeq | 7.2 |
| b3 | LEO | Subject3 | UEC741iPS517 | - | Nextera | 7.8 |
| b8 | LEO | Subject3 | UEC741iPS517 | + | Nextera | 7.1 |
| b9 | LEO | Subject3 | UEC741iPS517 | + | Nextera | 7.4 |
| b15 | LEO | Subject4 | HDF410iPS504 | - | Pilot-TruSeq? | 7.0 |
| b16 | LEO | Subject4 | HDF410iPS504 | - | Nextera | 7.4 |
| b17 | LEO | Subject4 | HDF410iPS504 | - | Nextera | 7.4 |
| b18 | LEO | Subject4 | HDF410iPS504 | - | Nextera | 7.7 |
| b25 | LEO | Subject4 | HDF410iPS504 | + | Nextera | 7.6 |
| b26 | LEO | Subject4 | HDF410iPS504 | + | Nextera | 7.7 |
| a8 | Ground | Subject3 | UEC741iPS517 | + | Nextera | 7.5 |
| a9 | Ground | Subject3 | UEC741iPS517 | + | Nextera | 7.6 |
| a13 | Ground | Subject4 | HDF410iPS504 | - | Nextera | 7.4 |
| a14 | Ground | Subject4 | HDF410iPS504 | - | Pilot-TruSeq | 7.2 |
| a15 | Ground | Subject4 | HDF410iPS504 | - | Nextera | 7.4 |
| a16 | Ground | Subject4 | HDF410iPS504 | - | Nextera | 7.9 |
| a23 | Ground | Subject4 | HDF410iPS504 | + | Pilot-TruSeq | 7.3 |
| a24 | Ground | Subject4 | HDF410iPS504 | + | Nextera | 7.4 |
| a25 | Ground | Subject4 | HDF410iPS504 | + | Nextera | 7.1 |

**Table S4**. Transcription Factor Enrichment Analysis

| **Cortical organoids: Transcription factor enrichment : Genes that are higher in LEO than ground** | | | **Cortical organoids: Transcription factor enrichment : Genes that are lower in LEO than ground** | | |
|---|---|---|---|---|---|
| **Rank** | **Transcription Factor** | **Hypergeometric p-value** | **Rank** | **Transcription Factor** | **Hypergeometric p-value** |
| 1 | SALL4 | 0.003052 | 1 | CEBPD | 0.000002643 |
| 2 | NFE2L2 | 0.02373 | 2 | KLF4 | 0.00001281 |
| 3 | SUZ12 | 0.02696 | 3 | TCF3 | 0.0002542 |
| 4 | REST | 0.03635 | 4 | NELFE | 0.0002565 |
| 5 | ESR1 | 0.03924 | 5 | SALL4 | 0.0003687 |
| 6 | FOXA1 | 0.0519 | 6 | BCL3 | 0.0005716 |
| 7 | SUZ12 | 0.06062 | 7 | ZBTB7A | 0.002032 |
| 8 | NFIC | 0.07004 | 8 | SUZ12 | 0.003086 |
| 9 | FOXA2 | 0.07901 | 9 | RFX5 | 0.004793 |
| 10 | BHLHE40 | 0.08672 | 10 | UBTF | 0.006349 |
| 11 | REST | 0.09509 | 11 | HNF4A | 0.007056 |
| 12 | RELA | 0.1189 | 12 | GATA2 | 0.007825 |
| 13 | EGR1 | 0.1313 | 13 | SIN3A | 0.008691 |
| 14 | MYC | 0.1396 | 14 | TAF7 | 0.009739 |
| 15 | TCF7L2 | 0.1418 | 15 | GATA1 | 0.01007 |
| 16 | SP1 | 0.1699 | 16 | TCF3 | 0.01195 |
| 17 | ZNF384 | 0.175 | 17 | CEBPB | 0.01728 |
| 18 | GATA2 | 0.1843 | 18 | ZMIZ1 | 0.01994 |
| 19 | SOX2 | 0.185 | 19 | ESR1 | 0.02054 |
| 20 | GATA1 | 0.192 | 20 | RUNX1 | 0.02074 |
| 21 | TCF3 | 0.1992 | 21 | NFIC | 0.02083 |
| 22 | E2F1 | 0.2034 | 22 | E2F6 | 0.02355 |
| 23 | AR | 0.2533 | 23 | MYOD1 | 0.02487 |
| 24 | SIN3A | 0.2607 | 24 | SOX2 | 0.02509 |
| 25 | SMC3 | 0.2709 | 25 | ATF2 | 0.02836 |
| 26 | RAD21 | 0.2878 | 26 | HDAC2 | 0.02842 |
| 27 | PBX3 | 0.2886 | 27 | FOXA2 | 0.03088 |
| 28 | RUNX1 | 0.2935 | 28 | TP53 | 0.0318 |
| 29 | TP63 | 0.3141 | 29 | PML | 0.03248 |
| 30 | MYC | 0.3363 | 30 | RELA | 0.03472 |

| | | |
|---|---|---|
| 31 | CTCF | 0.3868 |
| 32 | MAX | 0.4356 |
| 33 | ZBTB7A | 0.454 |
| 34 | CREB1 | 0.4627 |
| 35 | NFYA | 0.4647 |
| 36 | ATF2 | 0.5551 |
| 37 | E2F6 | 0.6076 |
| 38 | NFYB | 0.6642 |

| | | |
|---|---|---|
| 31 | FOSL2 | 0.03769 |
| 32 | CREB1 | 0.04007 |
| 33 | E2F1 | 0.04043 |
| 34 | TRIM28 | 0.04463 |
| 35 | PBX3 | 0.04484 |
| 36 | PPARG | 0.0496 |
| 37 | EGR1 | 0.05025 |
| 38 | ZNF384 | 0.05456 |
| 39 | CTCF | 0.0651 |
| 40 | NANOG | 0.07133 |
| 41 | SMC3 | 0.07135 |
| 42 | TRIM28 | 0.07539 |
| 43 | CHD1 | 0.09758 |
| 44 | ETS1 | 0.1166 |
| 45 | E2F4 | 0.1254 |
| 46 | NFYB | 0.1319 |
| 47 | STAT3 | 0.1336 |
| 48 | BHLHE40 | 0.1415 |
| 49 | STAT3 | 0.1427 |
| 50 | TP63 | 0.1477 |
| 51 | USF1 | 0.158 |
| 52 | TCF7L2 | 0.1732 |
| 53 | FOXA1 | 0.1793 |
| 54 | BCLAF1 | 0.2111 |
| 55 | FOS | 0.2141 |
| 56 | MAX | 0.2393 |
| 57 | VDR | 0.2496 |
| 58 | TAF1 | 0.2534 |
| 59 | IRF1 | 0.2583 |
| 60 | RCOR1 | 0.2665 |
| 61 | SP1 | 0.2707 |
| 62 | PPARD | 0.2886 |
| 63 | USF2 | 0.2912 |
| 64 | FOXM1 | 0.3011 |
| 65 | EGR1 | 0.3298 |
| 66 | NFE2L2 | 0.3333 |

| | | |
|---|---|---|
| 67 | MYC | 0.3347 |
| 68 | YY1 | 0.3529 |
| 69 | SPI1 | 0.3588 |
| 70 | ZKSCAN1 | 0.3636 |
| 71 | IRF8 | 0.3659 |
| 72 | ZC3H11A | 0.3847 |
| 73 | IRF3 | 0.4521 |
| 74 | NFYA | 0.4656 |
| 75 | FOXP2 | 0.4882 |
| 76 | REST | 0.5248 |
| 77 | BRCA1 | 0.5513 |
| 78 | STAT5A | 0.584 |
| 79 | EZH2 | 0.5902 |
| 80 | POU5F1 | 0.6258 |
| 81 | SMAD4 | 0.6456 |
| 82 | SRF | 0.676 |
| 83 | SP2 | 0.7249 |
| 84 | REST | 0.7646 |
| 85 | SPI1 | 0.8381 |
| 86 | NRF1 | 0.8454 |
| 87 | CREB1 | 0.8581 |
| 88 | RAD21 | 0.8603 |
| 89 | MYC | 0.8865 |
| 90 | YY1 | 0.9075 |
| 91 | AR | 0.9216 |
| 92 | ELF1 | 0.9995 |
| 93 | GABPA | 0.9998 |

| **Dopaminergic organoids: Transcription factor enrichment : Genes that are higher in LEO than ground** | | |
|---|---|---|
| **Rank** | **Transcription Factor** | **Hypergeometric p-value** |
| 1 | REST | 0.004137 |
| 2 | REST | 0.01957 |
| 3 | SUZ12 | 0.06471 |
| 4 | TCF3 | 0.1113 |
| 5 | AR | 0.1284 |
| 6 | EZH2 | 0.1798 |
| 7 | NANOG | 0.2963 |
| 8 | GATA2 | 0.3677 |
| 9 | KLF4 | 0.4458 |
| 10 | NFE2L2 | 0.4576 |
| 11 | RAD21 | 0.5339 |
| 12 | UBTF | 0.6309 |
| 13 | E2F6 | 0.8781 |
| 14 | TAF1 | 0.8868 |

| **Dopaminergic organoids: Transcription factor enrichment : Genes that are lower in LEO than ground** | | |
|---|---|---|
| **Rank** | **Transcription Factor** | **Hypergeometric p-value** |
| 1 | RELA | 5.060e-7 |
| 2 | AR | 0.00002807 |
| 3 | CEBPB | 0.0001756 |
| 4 | FOSL2 | 0.0008571 |
| 5 | TP63 | 0.001605 |
| 6 | ESR1 | 0.001827 |
| 7 | SALL4 | 0.003759 |
| 8 | NFIC | 0.004819 |
| 9 | TRIM28 | 0.006616 |
| 10 | TP53 | 0.008973 |
| 11 | NELFE | 0.01019 |
| 12 | MYOD1 | 0.01433 |
| 13 | SMAD4 | 0.016 |
| 14 | HDAC2 | 0.017 |
| 15 | STAT3 | 0.01764 |
| 16 | STAT3 | 0.01923 |
| 17 | PPARD | 0.02177 |
| 18 | TCF3 | 0.02317 |
| 19 | NFE2L2 | 0.02564 |
| 20 | TRIM28 | 0.03026 |
| 21 | FOXA2 | 0.03186 |
| 22 | TCF3 | 0.04318 |
| 23 | UBTF | 0.06425 |
| 24 | SUZ12 | 0.0791 |
| 25 | HNF4A | 0.09398 |
| 26 | RCOR1 | 0.09476 |
| 27 | FOXA1 | 0.1067 |
| 28 | CEBPD | 0.1127 |
| 29 | CTCF | 0.1152 |
| 30 | SMC3 | 0.1174 |
| 31 | PML | 0.1697 |
| 32 | RUNX1 | 0.1776 |
| 33 | IRF1 | 0.1801 |
| 34 | TCF7L2 | 0.2219 |

| | | |
|---|---|---|
| 35 | ZMIZ1 | 0.2422 |
| 36 | GATA2 | 0.2558 |
| 37 | SOX2 | 0.2586 |
| 38 | FOXP2 | 0.2602 |
| 39 | FOS | 0.2777 |
| 40 | TAF7 | 0.2809 |
| 41 | BHLHE40 | 0.3038 |
| 42 | VDR | 0.3465 |
| 43 | EGR1 | 0.3466 |
| 44 | CBX3 | 0.3643 |
| 45 | SPI1 | 0.3673 |
| 46 | ZNF384 | 0.379 |
| 47 | KAT2A | 0.3953 |
| 48 | RAD21 | 0.4006 |
| 49 | CREB1 | 0.4052 |
| 50 | NANOG | 0.4193 |
| 51 | POU5F1 | 0.4254 |
| 52 | ZBTB7A | 0.433 |
| 53 | SIN3A | 0.4361 |
| 54 | SUZ12 | 0.4435 |
| 55 | USF2 | 0.4449 |
| 56 | BCL3 | 0.4479 |
| 57 | ATF3 | 0.4706 |
| 58 | SPI1 | 0.4825 |
| 59 | IRF8 | 0.4908 |
| 60 | CHD1 | 0.4926 |
| 61 | SRF | 0.495 |
| 62 | E2F1 | 0.5186 |
| 63 | EGR1 | 0.5227 |
| 64 | USF1 | 0.5342 |
| 65 | PPARG | 0.5715 |
| 66 | KLF4 | 0.6431 |
| 67 | BRCA1 | 0.7225 |
| 68 | STAT5A | 0.7273 |
| 69 | EZH2 | 0.7333 |
| 70 | E2F6 | 0.7354 |
| 71 | NFYB | 0.7751 |
| 72 | RFX5 | 0.8191 |
| 73 | MAX | 0.8247 |
